# Nanovirseq: dsRNA sequencing for plant virus and viroid detection by Nanopore sequencing

**DOI:** 10.1101/2023.01.18.524564

**Authors:** Vahid J. Javaran, Abdonaser Poursalavati, Pierre Lemoyne, Dave T. Ste-Croix, Petter Moffett, Mamadou L. Fall

**Affiliations:** Saint-Jean-sur-Richelieu Research and Development Centre, Agriculture and Agri-Food Canada, Saint-Jean-sur-Richelieu, QC J3B 3E6, Canada; Département de Biologie, Centre SÈVE, Université de Sherbrooke, Sherbrooke, QC J1K 2R1, Canada; Département de phytologie, Faculté des Sciences de l’Agriculture et de l’Alimentation, Université Laval, QC, Canada, G1V 0A6

**Keywords:** dsRNAs, total RNAs, second- and third-generation sequencing technologies, grapevine virus and viroid detection, nanopore direct-cDNA and RNA sequencing, Illumina MiSeq sequencing

## Abstract

Worldwide, there is a need for certified clean plant materials to limit viral diseases spread. In order to design a robust and proactive viral-like disease certification, diagnostics, and management program, it is essential to have a fast, inexpensive, and user-friendly tool. The purpose of this study was to determine whether dsRNA-based nanopore sequencing can be a reliable method for the detection of viruses and viroids in grapevines or not. Compared to direct RNA sequencing from rRNA-depleted total RNA (rdTotalRNA), direct-cDNA sequencing from dsRNA (dsRNAcD) yielded more viral reads and detected all grapevine viruses and viroids detected using Illumina MiSeq sequencing (dsRNA-MiSeq). With dsRNAcD sequencing it was possible to detect low abundance viruses (e.g., Grapevine red globe virus) where rdTotalRNA sequencing failed to detect them. Indeed, even after removing rRNA, rdTotalRNA sequencing yielded low viral read numbers. rdTotalRNA sequencing was not sensitive enough to detect all the viruses detected by dsRNA-MiSeq. In addition, there was a false positive identification of a viroid in the rdTotalRNA sequencing that was due to misannotation of a host-driven read. For quick and accurate reads classification, two different taxonomical classification workflows based on protein and nucleotide homology were evaluated in this study, namely DIAMOND&MEGAND (DIA&MEG) and Centrifuge&Recentrifuge (Cent&Rec), respectively. Virome profiles from both workflows were similar except for grapevine endophyte endornavirus (GEEV), which was only detected using DIA&MEG. However, because DIA&MEG’s classification is based on protein homology, it cannot detect viroid infection despite giving more robust results. Even though Cent&Rec’s virus and viroid detection workflow was faster (30 minutes) than DIA&MEG’s (two hours), it could not provide the details and information DIA&MEG was able to provide. As demonstrated in our study, nanopore dsRNAcD sequencing and the proposed data analysis workflows are suitable and reliable for viruses and viroids detection, especially in grapevine where viral mixed infection is common.

## Introduction

The economic impact of grapevine production is of major importance in many countries, especially when it comes to wine production. In 2021, more than 7.3 million hectares of vineyards of various varieties of grapevines were planted in the world (1). Over the past few years, the economic potential of the grape and wine industry in Canada has expanded, both nationally and internationally. In 2019 in Canada, there were almost 12,545 hectares of vineyards planted, and the full economic impact of wine and grapes was over $11.5 billion (2). On the international scale, in 2021, both the volume and the value of exports from Canada have shown the greatest positive variations among major exporters. The volume of Canadian exports increased by 26% over 2020 to 2.1 million hectoliters. The bulk wine sector accounts for 99% of Canadian exports by volume and 68% by value (1). Nevertheless, increased outbreaks of viral and viroid diseases, which reduce grapevine growth, yields, quality of the fruit, and lifespan of the vineyards, represent a serious threat to the grapevine industry (3). A total of 89 viruses from 17 families and 34 genera, along with seven viroids from one family (*Pospiviroidae*) and four genera have been identified in infected grapevines (4-6). The four major viral diseases caused by around half of these identified viruses are infectious degeneration and decline, rugose wood complex, leafroll, and fleck disease (3, 5). Furthermore, the utilization of high-throughput sequencing (HTS) technologies has enabled all new viral species and new strains to be identified and detected, such as *Grapevine red blotch virus* (7-9), *Grapevine Syrah virus-1* (10), and *Grapevine Pinot gris virus* (11).

In the absence of effective chemical compounds for controlling viral diseases, the management of grapevine viruses is challenging (12). In addition to being able to adapt to various environmental situations and new hosts, viruses can evolve rapidly through mutation, genetic drifts, and genetic recombination (13). A number of other factors, including long periods of continuous monoculture, climate change, global trade of plant materials, and the expansion of the geographical range of insect vectors, have also led to the intensification of viral diseases (14-17). Consequently, growers need to identify viruses as early as possible in order to take timely action and implement necessary sanitary measures (4, 5, 18).

Even though several advanced and traditional diagnostic methods are available for grapevine viruses, including immunological (19), nucleic acid amplification (20), microarrays (21), and hyperspectral imaging (22, 23), their inability to detect all known viruses simultaneously as well as novel and unknown viruses is still one of their major limitations. The introduction of second-generation of sequencing (SGS) has resulted in the detection and identification of many novel and known grapevine viruses, including grapevine Syrah virus 1 (GSV1), grapevine vein clearing virus (GVCV), grapevine pinot gris virus (GPGV), grapevine virus F (GVF), grapevine red blotch virus (GRBV), grapevine roditis leaf discoloration-associated virus (GRLDaV), grapevine virus N (GVN) and grapevine virus O (GVO) (6, 7, 10, 24-26). Although SGS has detected and discovered known and unknown viruses and been used as a high potential diagnostic tool, its limitations have slowed its usage compared to other methods in diagnostic labs. These include, laborious and expensive library preparation techniques, data management, expensive sequencing equipment, and the need for sophisticated technical expertise in order to analyze data (27-30). Further, in routine diagnostic laboratories, small number of samples may need to be sequenced, and using SGS would not be economically viable (31). Some of these limitations have been addressed with the introduction of third-generation of sequencing (e.g. nanopore sequencing technology) (4, 32, 33).

A number of features, such as the small size of the sequencer (MinION), the ease of library preparation, the low sequencing cost, the possibility of long-read sequencing, and the rapid speed of the sequencing process, make it an excellent tool for the surveillance of viruses and other pathogens (4, 34, 35). Plant virus detection has been conducted using various DNA and RNA nanopore sequencing kits, and this sequencing technology has shown potential for diagnostic purposes. Since a majority of the plant viruses are RNA viruses, cDNA and native RNA-based kits, such as direct RNA sequencing, direct cDNA sequencing, and cDNA-PCR sequencing, are frequently used (4, 35). As RNA viruses usually do not contain poly A tails, in order to sequence them via nanopore sequencing kits that use poly T adapters, a number of modifications need to be made during the library preparation steps. The following options are available for sequencing poly A tailed and non-poly A tailed viruses: a random hexamer primer strategy for cDNA synthesis, which can be used only for cDNA sequencing kits and adding several adenine nucleotides to the 3’ end of RNA with E. coli poly A polymerase (35). In addition to RNA viruses, several DNA viruses (single or double stranded) have been detected by nanopore sequencing technology. For instance, the nanopore sequencing rapid barcoding kit, which can be used in the field, was able to detect African cassava mosaic virus and East African cassava mosaic virus (36).

Despite the fact that different nucleic acid types (DNA or RNA) and various library preparation strategies have been used for plant virus detection by nanopore sequencing (31, 36-44), there is only two reports of plant virus detection through double-stranded RNA (dsRNA) nanopore sequencing and there were related to single virus infections by new isolates of *Jasmine Virus C* (45) and cucumber Bulgarian latent virus (46). Plant virus detection is usually accomplished using total RNA (31, 47) but using total RNA has a number of limitations. The majority of total RNA sequencing reads are related to host transcripts, such as ribosomal RNAs and messenger RNAs. In order to achieve more specific diagnostic results, undesirable RNA must be removed prior to library preparation, which is costly. A good alternative for detecting plant viruses is to use dsRNA, an intermediate replicative RNAs molecule generated during the virus transcription process (48, 49). Even though it was not initially confirmed that negative-sense single-stranded RNA viruses (-ssRNA) produce dsRNAs in their replication (50), recent viromics studies have shown that these viruses also produce dsRNAs in small amounts (51-53). The use of dsRNA in our previous research also enabled the detection of not only RNA viruses and viroids, but also of the grapevine red blotch-associated virus (GRBV), a DNA virus (24). As a result, dsRNA is a suitable starting material for the detection of viruses regardless of their genomic materials.

The aim of this study was to introduce a simple nanopore dsRNA (dsRNAcD) sequencing protocol, not only for diagnostic laboratories, but also for evolutionary studies. We describe step-by-step a protocol that can be used to support diagnostic testing of infected grapevine samples using the Oxford Nanopore Technologies (ONT) MinION sequencing device. Grapevine was selected as a challenging plant that is host to multiple viruses with many virus mixing infection events and nucleic acids extraction inhibitors. The dsRNA extraction protocols and library preparation protocols for grapevine samples were optimized by taking into account the sequencing price per sample. In these experiments, direct RNA sequencing and direct cDNA sequencing kits were used for library preparation, and the performance of each kit for detecting viruses in various experimental circumstances was tested. Also, the results were compared to those from Illumina sequencing in order to draw comparative conclusions between the two sequencing technologies when it comes to detecting viruses. Moreover, a cost-effectiveness analysis was completed to determine when it is suitable to use this technology. Finally, two different bioinformatics workflows depending for the diagnostic or evolutionary purposes were presented. Overall, dsRNAcD sequencing has great potential for plant virus and viroid detection and genomic characterization in case of mixed infections. dsRNAcD sequencing can greatly reduce sequencing costs. Multiple samples can be sequenced on the same flow cell at the same time, which could lead to substantial cost savings compare to SGS.

## Material and methods

### Plant material

A total of 24 asymptomatic and symptomatic grapevine samples (a combination of leaves and petioles) were collected from a vineyard at Agriculture and Agri-Food Canada’s experimental farm in Frelighsburg (latitude 45°03′12′′ N; longitude 72°51′42′′ W), Quebec (Supplementary file S1). Samples were collected from grapevine (*Vitis vinifera* ‘Vidal blanc’) plants over the course of July and September 2019 and placed into sterile 50-mL centrifuge tubes for cold storage at 20°C. The leaves were washed with distilled water, roughly crushed, and homogenized in a liquid nitrogen-cooled 50-mL conical centrifuge tubes with 8 stainless-steel balls (8 mm) via a 1600 MiniG® Tissue Homogenizer and Cell Lyser (Spex® SamplePrep). Next, powdered leaves (1.5-2 g) were transferred into sterile 50-mL centrifuge tubes and stored at -80°C until they were ready for the nucleic acid extraction.

### dsRNA extraction

An adapted version of Fall et al. (24) and Kesanakurti et al. (54) dsRNA extraction protocols was used to extract dsRNA from 24 different grapevine samples In brief, 12 ml of extraction buffer (200 mM Tris (pH 8.3), 10 mM EDTA, 300 mM Lithium chloride, 55 mM lithium dodecyl sulfate, 25 mM deoxycholic acid, 2% PVP-40000, 1% Nonidet P-40, and 1% 2-Mercaptoethanol) were added to 1.5 g homogenized leaf samples. In addition, a positive control, *Phaseolus vulgaris* cultivar Black Turtle Soup (BTS) known to be infected by *endornavirus 1* (PvEV1) and *Phaseolus vulgaris endornavirus 2* (PvEV2) (24, 54), was added at a final concentration of 1% (w/w) in each samples to assess the efficiency of the dsRNA extraction protocol. After 40 minutes of shaking at 300 rpm, the tubes were centrifuged at 1000 x g for 1 minute at 10°C to remove bubbles and debris. As soon as the supernatant was transferred into a new 50-ml tube, 12 ml of potassium acetate buffer (5.8 M) was added, and the tubes were centrifuged at 14,000 x g for 15 minutes at 10°C. After transferring the supernatant into a clean 50-ml centrifuge tube, 16 ml of 100% isopropanol was added, and the tubes were stored at -20°C for 20 minutes. Centrifugation was performed at 11,000 x g for 16 minutes at 4°C, then the supernatant was discarded, and the pellet was dissolved in STE-18 (10 mM Tris (pH 8.0), 100 mM NaCl, 1 mM EDTA (pH 8.0), and 18% ethanol). Next, we add 300 mg of Sigmacell Cellulose Type 101 dissolved in 2 ml of STE-18 to the solution. the tubes were shaken at 300 rpm for 15 minutes at room temperature, centrifuged at 14,000 x g for 5 minutes, and then the supernatant was discarded. To discard single-strand RNAs and DNAs, we performed two washing steps using STE-18, the first with 40 ml and the second with 20 ml. The supernatant was discarded by centrifuging at 14,000 x g for 5 minutes at 20°C between washing steps. Finally, to elute the extracted dsRNA, 6 ml of 1XSTE (10 mM Tris (pH 8.0), 1 mM EDTA (pH 8.0), and 100 mM NaCl) was added to the cellulose pellet and the solution was stirred for 15 minutes on the shaker. After centrifugation at 14,000 x g for 8 minutes at 20°C, the supernatant was transferred to a new 50-ml centrifuge tube, and 3M sodium acetate (pH 5.2) and alcohol were used to precipitate dsRNAs. The detailed protocol can be found in the protocols.io website (https://www.protocols.io/view/double-stranded-rna-extraction-by-cellulose-4r3l2odrjv1y/v1).

### Total RNA extraction

Three different samples randomly selected among the 24 samples collected. Based on the MacKenzie et al. (55) protocol, total RNAs were extracted from 100-mg leaf material of them, using the RNeasy Plant mini kit (Qiagen, Canada). Quantitative and qualitative measurements of total RNA were performed using a NanoDrop 2000c (Thermo Sci, Canada) and a Qubit 4 Fluorometer (Life Technologies, Canada).

### dsRNAcD sequencing library preparation

To ensure complete removal of ssRNAs and DNAs, dsRNA were digested with DNase I and RNase T1. Digestion was stopped by adding EDTA 50 mM and heating at 65°C for 10 minutes. In the presence of 2 µl of 60 µm random primers, 1 µl of 10 mM dNTP, and 6 µl of water, dsRNA was denatured at 99°C for five minutes. Then tubes were immediately placed in ice water and a master mix (4 µl of first strand cDNA synthesis buffer, 1 µl of RNase out or RNasin® Ribonuclease Inhibitor (40 u/µl), and 1 µl (200 units) of Maxima H minus) was added. The reverse transcription step was performed for 90 min at 55°C. One unit of Ribonuclease H was then used to hydrolyze the DNA-RNA duplex. The second strand of cDNA was synthesized by adding Klenow DNA Polymerase I and *E. coli* DNA Ligase I. Agencourt AMPure XP magnetic beads (Beckman-Coulter) were used to clean up two-stranded cDNAs. The detailed protocol can be found in the protocols.io website (https://www.protocols.io/view/synthesis-of-double-strand-cdna-ds-cdna-from-viral-bp2l69nddlqe/v1).

With the direct cDNA sequencing kit (SQK-DCS109, ONT) and its protocols, two libraries of cDNA samples were generated. Initially, 24 cDNA samples from various infected grapevines were pooled to prepare a library (called pooled library) using the direct cDNA Sequencing protocol (DCS_9090_v109_revO_14Aug2019) without any multiplexing barcodes. Using direct cDNA sequencing kit with native barcoding (EXP-NBD104 and EXP-NBD114) kits, the second library (called multiplexed library) was prepared for 23 different cDNA samples according to the manufacturer’s recommendations (Figure 1); there was one sample from 24 collected samples failed to pass the library preparation steps. Since the cDNA synthesis was performed by random primers, the library preparation process was started from the “End-prep” step of the aforementioned protocol using NEBNext Ultra II End repair/dA-tailing Module (New England Biolabs,NEB). A Blunt/TA Ligase Master Mix (NEB) was then used to ligate the sequencing adapter (AMX) to the pooled library. The multiplexed library was constructed by ligating a native barcode to each sample using Blunt/TA Ligase Master Mix. In order to pool barcoded samples together equally, we measured the quantity of each sample with a Qubit dsDNA HS Assay Kit and a Qubit 4.0 fluorometer. As a final step, NEBNext Quick Ligation Module (NEB E6056) was used to ligate the sequencing adapter (AMII) to the multiplexed library. Following each enzymatic step of the protocol, AMPure XP magnetic beads were used to purify the samples. It should be mentioned that, following each enzymatic step of the protocols, AMPure XP magnetic beads were used to purify the samples.

**Figure 1.**
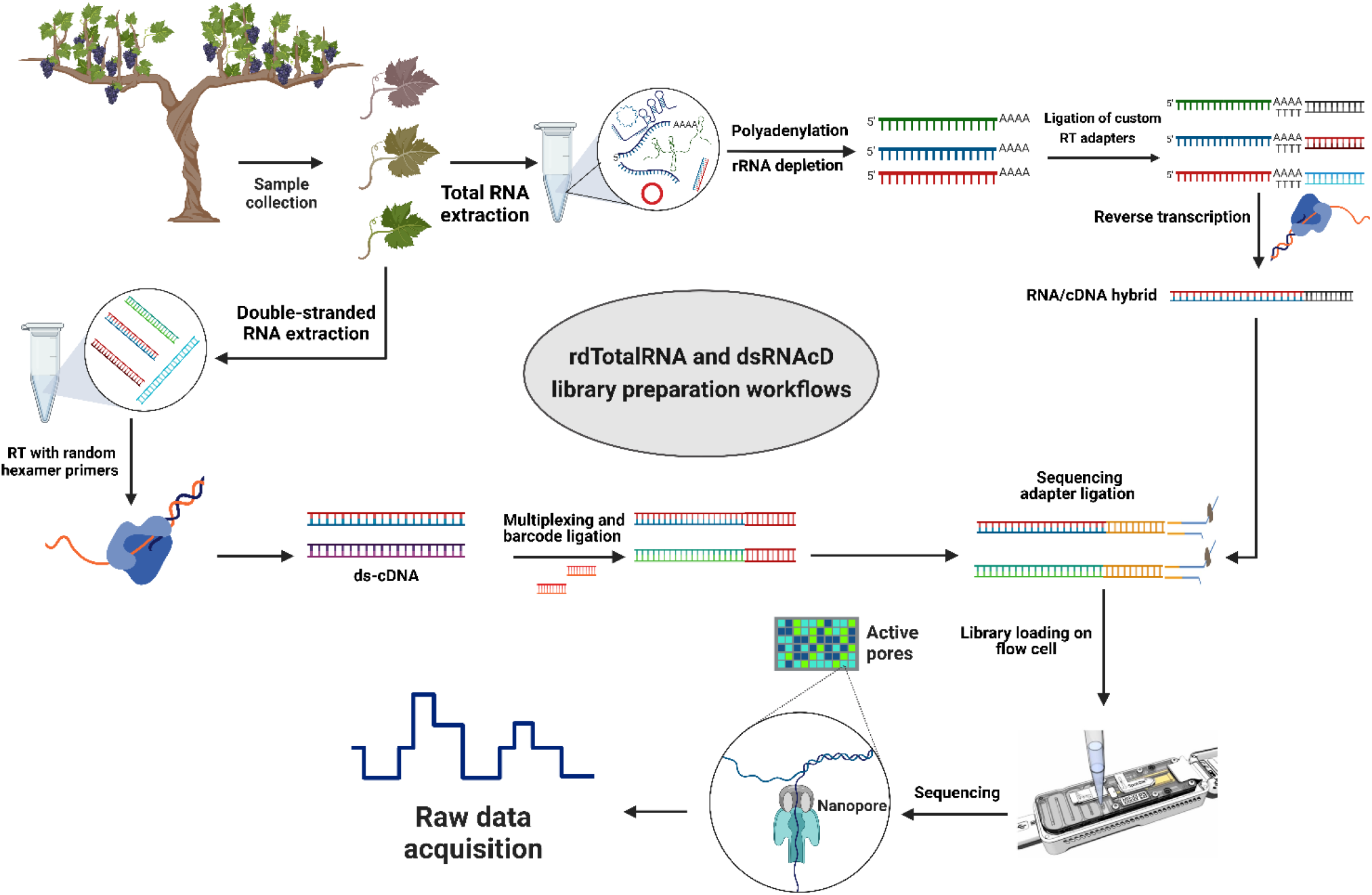
The library preparation workflows for grapevine virus and viroid detection. Direct RNA sequencing (top): Initially, rRNAs were depleted from total RNA and then polyadenylated. Next, different custom reverse transcription adapters were ligated to polyadenylated transcripts, and after reverse transcription (RT), the sequencing adapter was ligated, and a pooled library was loaded on a R9.4.1 flowcell. **Direct cDNA sequencing (left-down)**: Following dsRNA extraction, double-stranded cDNA was synthesized using random primers followed by Klenow polymerase and E. coli DNA Ligase to create second strand cDNA. A commercial barcode was ligated to each sample after double-stranded cDNAs were end prepared. The pooled library was made from 23 different barcoded samples, and after ligating the sequencing adapter, the pooled library was primed and loaded on another R9.4.1 flowcell. (The figure was designed by BioRender.com)

### rdTotalRNA sequencing library preparation

The direct RNA sequencing kit (SQK-RNA002) from ONT is optimized for poly A-tailed transcripts. Therefore, several modifications were made to the direct RNA sequencing library preparation protocol to capture both poly A-tailed and non-poly A-tailed viral RNAs. After DNase I digestion and removing the ribosomal RNAs (rRNAs) from total RNA using the QIAseq FastSelect-rRNA Plant Kit probe (QIAGEN), several Adenine bases were tailed at the 3’ end of the remaining RNAs according to Liefting et al (47). The samples were then multiplexed using three pre-annealed RT adaptors ordered from Integrated DNA Technologies (IDT), based on the DeePlexiCon method (Supplementary file S2) (56). In brief, each custom RT adapter was ligated to 500 ng of rRNA-depleted and polyA-tailed RNA samples using T4 DNA Ligase (NEB M0202L), which was followed by reverse transcription using SuperScript III Reverse Transcriptase (Thermo Fisher Scientific). Purification of cDNA/RNA hybrid complexes as reverse transcription products was achieved using Agencourt RNAClean XP beads. Qubit dsDNA HS Assay Kit was used to measure each sample’s concentration, and 65 ng of reverse transcribed RNA was taken from each sample to pool the samples together at equal concentrations. The RNA sequencing adapter (RMX), was ligated to the RNA-cDNA hybrid complex using T4 DNA Ligase (NEB M0202L) and subsequently purified with the Agencourt RNAClean XP beads (Beckman-Coulter) at a ratio of 1X according to the Direct RNA Sequencing protocol (Figure 1).

### Priming and loading the R9 Flow Cell, sequencing, demultiplexing, and basecalling

Three different nanopore sequencing libraries (Table 1) were loaded on three MinION R9.4.1. (FLO-MIN106D) flow cells by using the Flow Cell Priming Kit (EXP FLP002) according to the manufacturer’s instructions. The sequencing step was carried out on MinION Mk1B device, and the sequencing conditions were set up using MinKNOW software (v.21.11.8). Following 24 hours of sequencing and raw data acquisition, the raw data were basecalled and demultiplexed by Guppy software (v6.0.6). Score of seven was considered as the minimum quality score, and reads below this score were removed. In the direct RNA sequencing experiment, raw data were basecalled using Guppy software (v6.0.6), and demultiplexing was carried out using DeePlexiCon tool according to the developer’s instruction.

**Table 1.** A description of the samples, sequencing kits, and library preparation information used in this study.

|  | Number of samples | Sequencing kit | Material | Multiplexing |
| --- | --- | --- | --- | --- |
| **Library 1** | 24 | Direct cDNA sequencing<br>(SQK-DCS109) | dsRNA | None |
| **Library 2** | 23 | Direct cDNA sequencing<br>(SQK-DCS109) | dsRNA | Native Barcoding Expansion kits<sup>a</sup><br>(EXP-NBD104 and EXP-NBD114) |
| **Library 3** | 3 | Direct RNA sequencing<br>(SQK-RNA002) | Total RNA | Custom barcodes<sup>b</sup> |
| **Library 4** | 23<sup>c</sup> | Illumina Nextera XT DNA<br>library prep kit | dsRNA | Unique dual indexes |

- Native Barcoding Expansion kits contain 24 unique barcodes for multiplexing and sequencing different samples simultaneously on one flow cell. - Custom barcodes are pre-annealed RT adaptors which were designed based on DeePlexiCon method for multiplexing and sequencing different RNA samples simultaneously on one flow cell. - The dsRNA materials used for MiSeq sequencing were the same as those used for dsRNACD sequencing.

### Quality control and preprocessing of datasets

In order to analyze the quality and extract descriptive statistics from raw sequencing data of the three nanopore sequencing libraries, NanoPlot (version 1.33.0) (57) was used. To trim each dataset individually, separate quality plots from the head and tail regions of reads were depicted by NanoQC (v0.9.4). The head and tail of each read were then trimmed using NanoFilt (v2.8.0) (57). Quality control was repeated again by NanoPlot (version 1.33.0) to extract the statistics results. The host sequence contaminations were removed by aligning reads against Grapevine genome (GCF_000003745.3_12X) using Minimap2 software (v2.17-r941) (58) for nanopore sequencing datasets and bowtie2 (59) for MiSeq datasets and excluding reads related to the host by SAMtools (v1.6) (60) to increase data analysis speed and accuracy. Clustering, error correction, and polishing of the trimmed and filtered datasets were performed using Rattel toolbox (v1.0) (61) in accordance with the developer’s instructions for nanopore sequencing datasets. In the case of direct RNA sequencing, Rattle was run with the “*-y rna*” option.

### Illumina library preparation and sequencing

We synthesized 23 cDNA samples using the same dsRNA materials that were used for dsRNAcD library preparation, according to the cDNA synthesis procedure in dsRNAcD library preparation section. An Illumina Nextera XT DNA library prep kit was used to prepare the libraries from 1 ng of double-stranded cDNA input. A paired-end sequencing step with the MiSeq Reagent Nano Kits V2 was conducted using the Illumina MiSeq sequencer as described in Fall et al 2020 (24).

### Nanopore sequencing data analysis

#### Strategy 1: Virus and viroid detection through Centrifuge-Recentrifuge (Cent&Rec) workflow

After quality control, read trimming, cleaning and clustering etc. (preprocessing step in Figure 2), the long reads were classified taxonomically by Centrifuge classifier (version 1.0.4) using an in-house workflow with a customized Centrifuge indexing database (CID) (Figure 2) (62). The CID was constructed using a local database (2021), which included GenBank, RefSeq, TPA and PDB genome, gene and transcript sequence data of viruses, together with *Homo sapiens* GRCh38p13 assembly, bacteria, archaea and ViroidDB. First, taxonomic assignment was carried out by Centrifuge using default values. Second, Recentrifuge software (version 1.9.1) (63), was used for comparative analysis of the classification results and to produce interactive HTML reports using the *-y* 50 option (Figure 2). To further curate the taxonomic results, Blastn and Blastx (64) were also conducted in this section.

**Figure 2.**
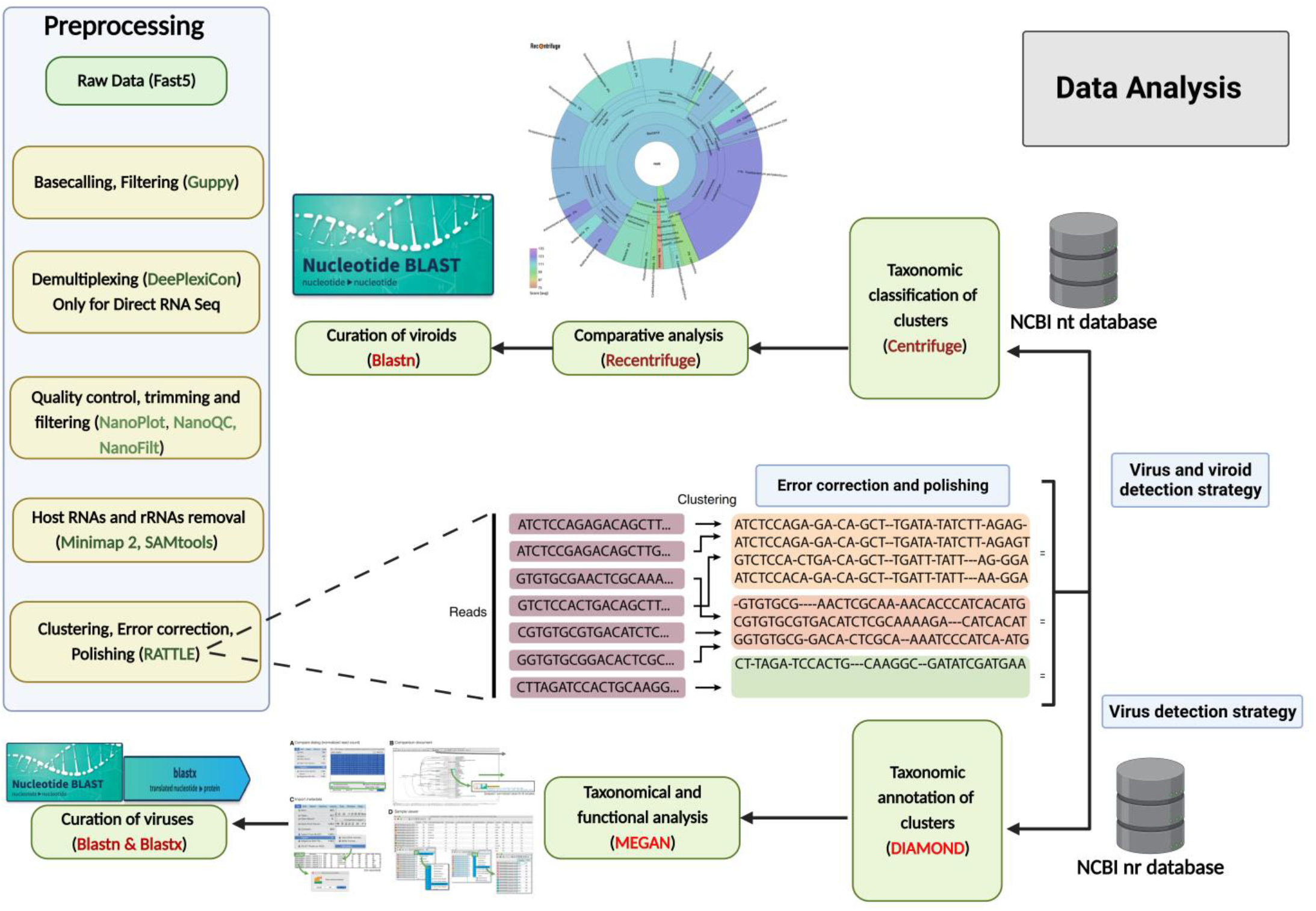
Data analysis workflows for grapevine virus and viroid detection through nanopore sequencing. Preprocessing of raw data (left): After the acquisition of raw data through MinKNOW software, basecalling, filtering according to quality score (<7), demultiplexing, trimming, removing host reads, clustering, correcting errors and polishing were conducted using different software and packages. **Centrifuge and Reecentrifuge strategy (right-top):** Pre-processed reads were taxonomically assigned by Centrifuge, and then comprehensive analysis was conducted and visualized by Recentrifuge. **DIAMOND+MEGAN strategy (right-down)**: In addition, the same reads from the previous step were aligned with annotated protein sequences (NCBI-nr) by DIAMOND, and subsequently, MEGAN 6 binned the sequences based on their taxonomy and function profiles. To further curate the taxonomic results, Blastn and Blastx were also conducted.

#### Strategy 2: Virus detection through DIAMOND-MEGAN workflow (DIA&MEG)

By aligning corrected and clustered reads against a database of annotated protein sequences (NCBI-nr), a taxonomically and functionally binning of the sequences were completed following the Bağcı et al. procedure with some minor modifications (65). Instead of processing long read datasets through a de novo assembly step as described in Bağcı et al. procedure, DIAMOND (66) was used directly to align the corrected reads against NCBI-nr without any de novo assembly step, and then the taxonomic and functional binning steps were completed with MEGAN 6 (67) with one modification. Instead of “LongReads” parameter, “weighted” parameter were selected in meganization step. Interactive analysis of the results was performed using MEGAN 6, and comparative bar charts and taxonomic trees were extracted from this software (Figure 2). To further curate the taxonomic results, Blastn and Blastx (64) were also conducted.

### Illumina MiSeq Data analysis

MiSeq sequencing short reads were analyzed in two ways: 1. Based on the LazyPipe pipeline (68), which in this manner initially short reads were de novo assembled, then taxonomic profiling was performed by the Centrifuge classifier, and finally, a comparative analysis was conducted by Recentrifuge. 2. The DIAMOND-MEGAN workflow described above were used to align the short reads against NCBI-nr and produce comparative bar charts and taxonomic trees.

## Results

### Library preparation and sequencing efficiency

The aim of this study was to develop and evaluate two types of nanopore sequencing strategies, rdTotalRNA sequencing and dsRNAcD sequencing, for detecting viruses and viroids in mixed-infected grapevine samples. The pooled dsRNA library from 24 different grapevine samples was sequenced using nanopore direct cDNA sequencing yielded a total of 1,754,037 reads after 24 hours. The 23 barcoded samples were sequenced on another flow cell which yielded 2,921,438 reads (Supplementary file S3). In addition, to compare dsRNAs and total RNAs as starting material, three out of the 23 aforementioned samples (CO-9-86J, BV-12-16J and BIO-15-56S) were also selected randomly and used for total RNA extraction and direct RNA sequencing library preparation. After sequencing of the rdTotalRNA library, the 541,972 reads which were produced were demultiplexed and basecalled (Supplementary file S3). The dsRNA-MiSeq sequencing also yielded 194,549, 340,946, and 460,909 sequences with quality scores above Q30 for CO-9-86J, BV-12-16J, and BIO-15-56S samples, respectively. For processing and sequencing the samples, rdTotalRNA sequencing (29.17 h) and dsRNAcD sequencing (37.58 h) were faster than Illumina dsRNA-MiSeq (88.72 h). Nanopore rdTotalRNA sequencing was 1.3 and 3 times faster than nanopore dsRNAcD sequencing and dsRNA-MiSeq sequencing, respectively. (Table 2). In terms of sequencing cost, nanopore dsRNAcD sequencing was significantly cheaper (Can$103 per sample) than nanopore rdTotalRNA sequencing (Can$350 per sample) and Illumina dsRNA-MiSeq (Can$412 per sample) (Supplementary file S4).

**Table 2.**
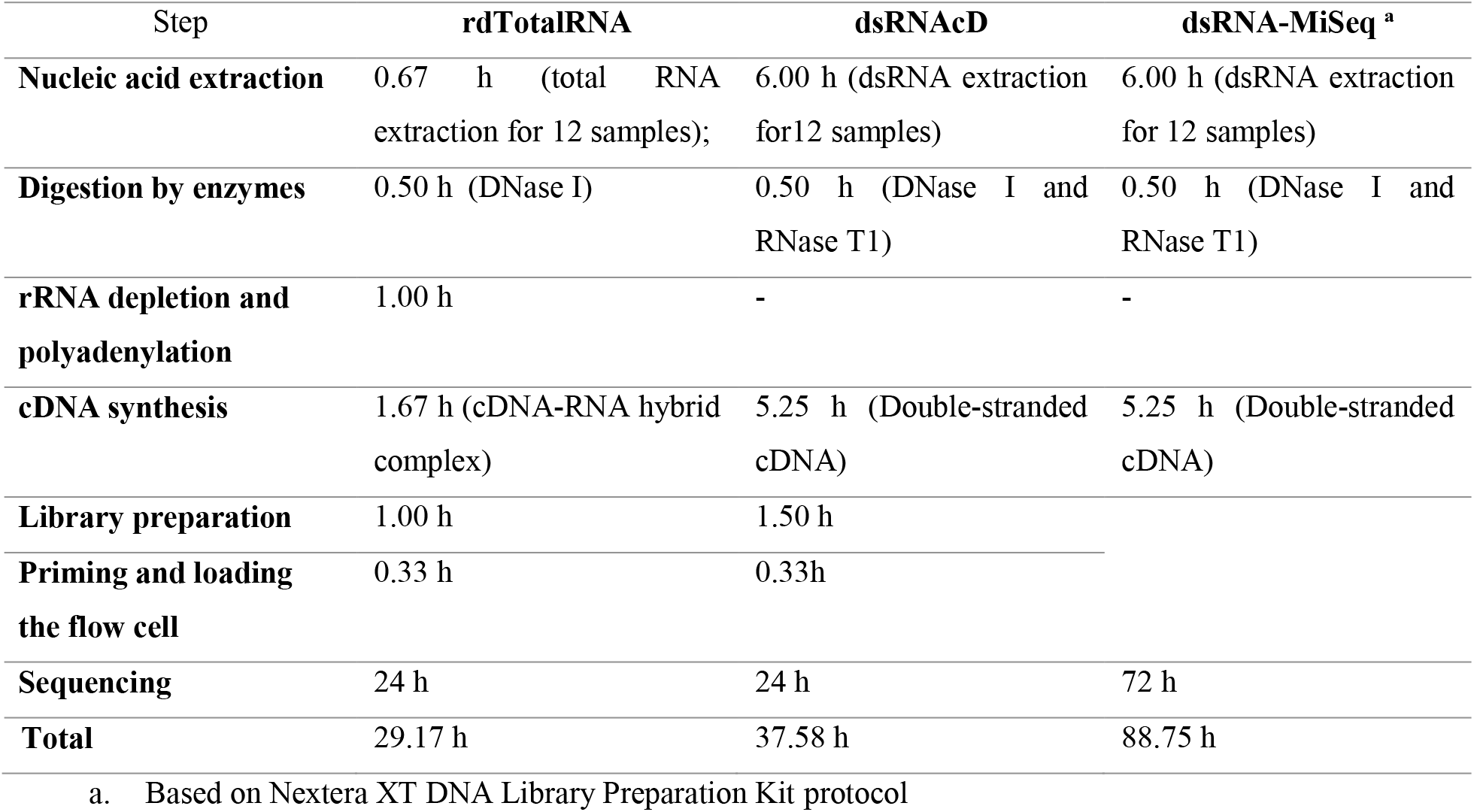
The estimated times for extraction, library preparation, and sequencing of each RNA extraction and sequencing method.

### Preprocessing, clustering and error correction of long reads

For the sake of convenience, among the 23 samples in this study which were sequenced using dsRNAcD sequencing and dsRNA-MiSeq, three samples were randomly chosen to be sequenced by rdTotalRNA as well. However, the results from the remaining 20 samples were presented in the supplementary materials. The statistics of raw data revealed that even though all barcoded samples were pooled together at the same concentrations, the number of sequenced reads varied between samples (Table 3). Nanopore sequencing protocols are optimized for long reads. Because several bead purification steps were carried out with different ratios during library preparation, samples containing short nucleic acid fragments lost more sequences, which resulted in fewer reads than those containing long fragments. The mean quality of reads was improved in all samples with the trimming and filtering processes (Table 3 and Supplementary file S3). In addition, removing unwanted host related reads from the rdTotalRNA datasets revealed that 47-67% of the raw reads were related to the host, while in the dsRNAcD datasets, host-related reads ranged from 21-65%. Before taxonomical classification, error correction and clustering of reads also increased the mean quality of corrected reads in both kinds of nanopore sequencing datasets (Table 3).

**Table 3.**
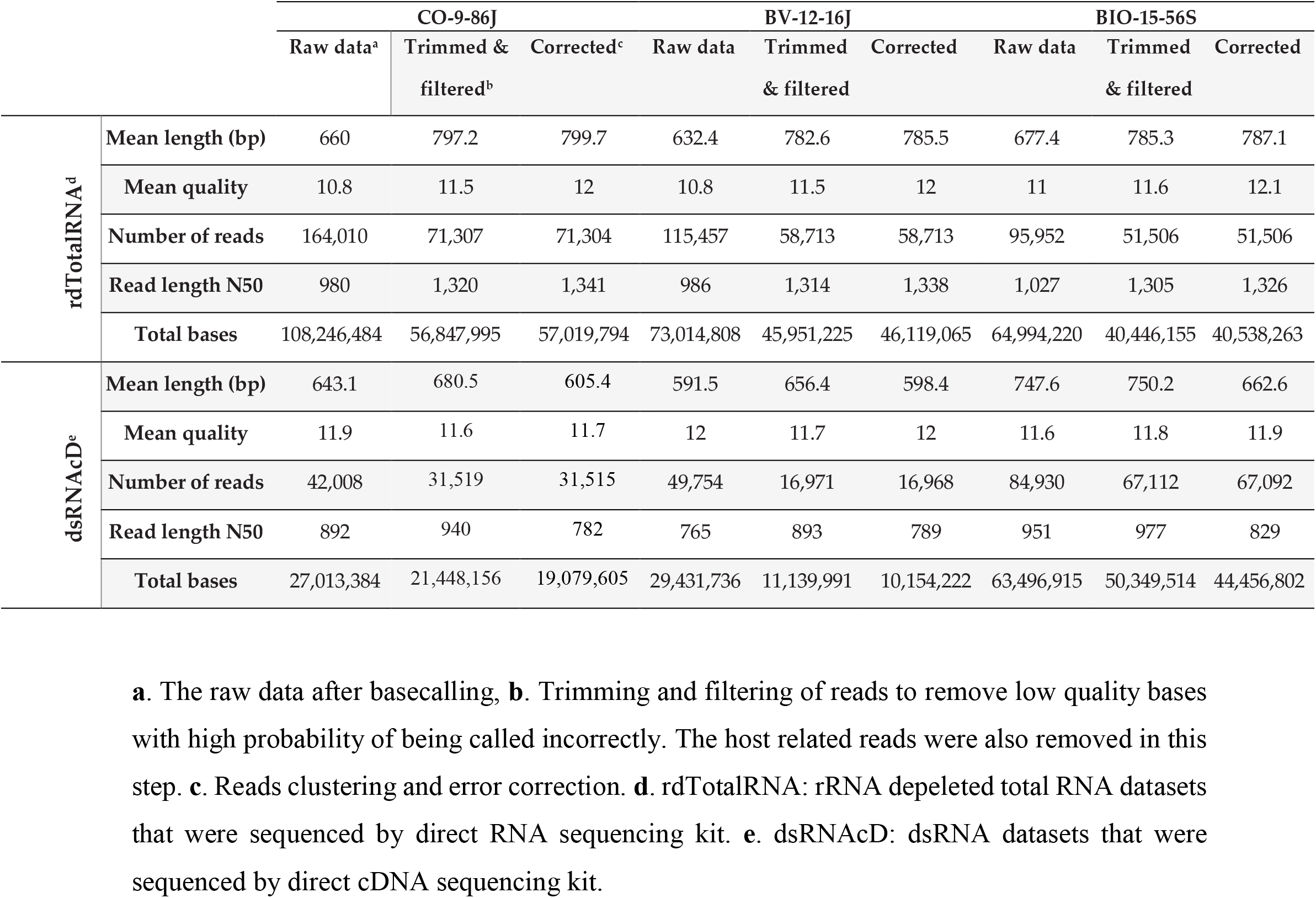
Overview of nanopore sequencing raw data preprocessing, filtering, trimming, clustering and error correction.

### Nanopore data analysis strategy 1: Centrifuge and Recentrifuge (Cent&Rec)

For both dsRNAcD and rdTotalRNA sequencing datasets, a customized index database which included complete genome sequences of all possible viruses was used for taxonomic classification by Centrifuge Classifier (62), a sensitive and high speed classifier designed to classify sequences and accurately process millions of reads within a few minutes (69). The frequency of reads that could not be taxonomically assigned to subject reads in our customized index database were different between dsRNAcD and rdTotalRNA libraries (Table 4); it was between 36.5 and 55% for dsRNAcD libraries and 1.5 to 16% for rdTotalRNA libraries. Some of these unassigned reads in dsRNAcD libraries may have been related to novel virus strains or species.

**Table 4.**
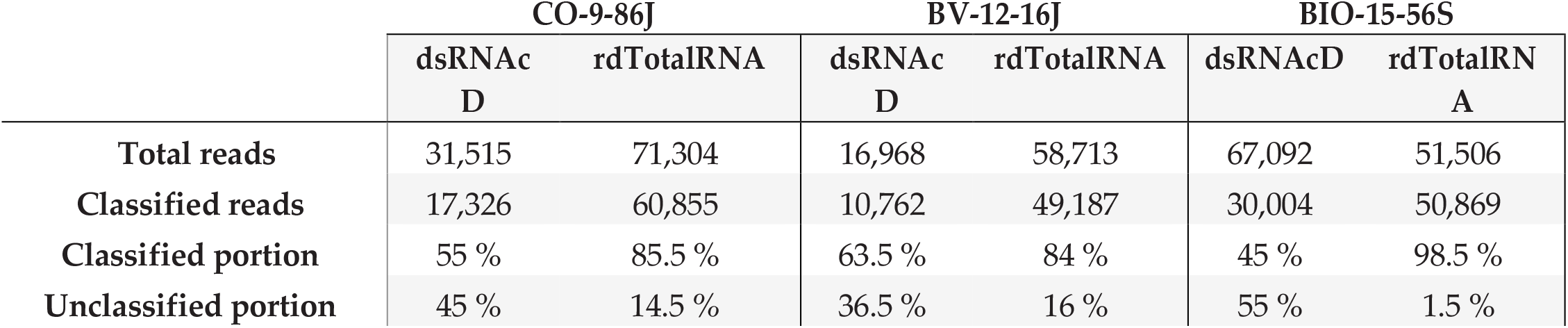
Centrifuge’s classification performance for dsRNAcD sequencing and rdTotalRNA sequencing datasets, with the classified and the unclassified portion (%) of reads.

While the sequencing platforms and analysis workflows were different for nanopore sequencing and MiSeq, we used the same index database for long and short dataset taxonomy classifications. In order to perform a comparative analysis of the results obtained during the taxonomical classification, Recentrifuge Software (63) was used, and both the score-based visual results and statistical results were extracted as HTML charts (Figure 3) and Excel files (Supplementary Folder 1). In all samples, reads derived from the host were initially subtracted for more accurate taxonomical classification and to increase viral read portions. In several studies, it was shown that the subtraction of host coextracted sequences from short and long read datasets reduced CPU time for taxonomical classifications and helped discover and characterize viral sequences present at low titers (70, 71). The viral portion of assigned reads in dsRNA libraries sequenced by dsRNAcD and dsRNA-MiSeq were significantly higher (in the range of 85% to 95%) than the viral portions in rdTotalRNA libraries (in the range of 6% to 21%) (Supplementary Folder 1).

**Figure 3.**
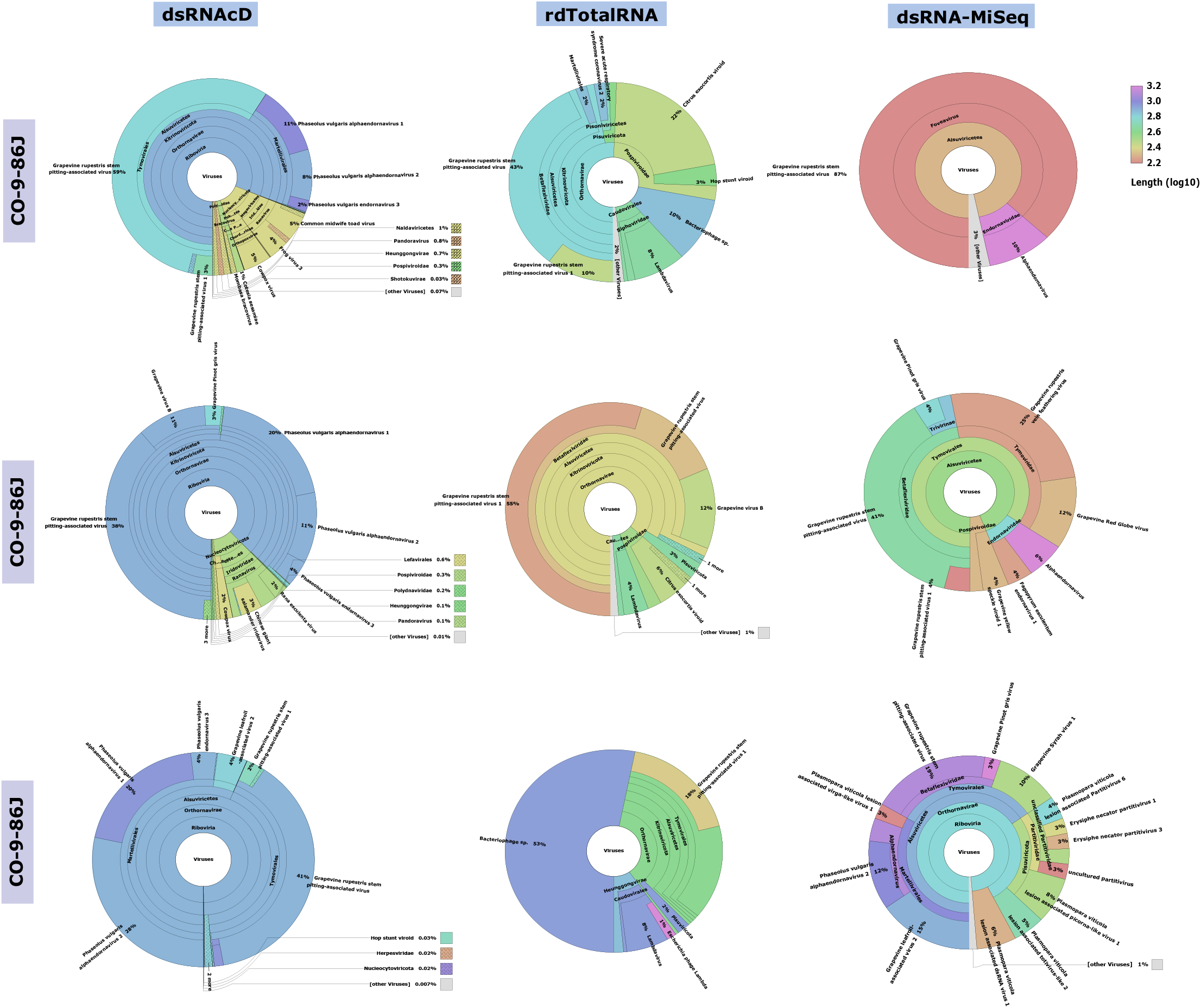
Comparative analysis of Centrifuge’s results by Recentrifuge software. Recentrifuge HTML interface screenshots for different samples and sequencing technologies are shown. The virus super kingdom is drawn in the center with its hierarchical pie chart zoomed in. Depending on the confidence level of the taxonomic classification, the background color varies for each taxon. For the results from 20 other samples, see supplementary folder 2.

In all dsRNA samples, the spiked positive control bean viruses (PvEV1, PvEV2, and PvEV3) were detected. The grapevine viruses detected by dsRNAcD and by dsRNA-MiSeq sequencing were same in most of the samples. In few samples, dsRNAcD was the only method for detecting viruses whose abundance was low in our samples, including GLRaV-2, GLRaV-3, and GRGV. In contrast, rdTotalRNA sequencing wasn’t able to detect most of viruses that were detected by dsRNAcD sequencing and dsRNA-MiSeq (Table 4). The rdTotalRNA sequencing performed better with viruses that were highly abundant (based number reads) in a sample such GRSPaV and GVB from the Betaflexiviridae family. These results suggested that nanopore dsRNAcD sequencing was more sensitive to low abundance viruses than rdTotalRNA sequencing and produced similar results compare to dsRNA-MiSeq. In addition to the three samples presented in this section (Figures 3 and 4), 20 other samples analyzed using dsRNA-MiSeq and dsRNAcD sequencing also provided similar results (Supplementary Folder 2) demonstrating the repeatability and accuracy of the dsRNAcD sequencing method.

**Figure 4.**
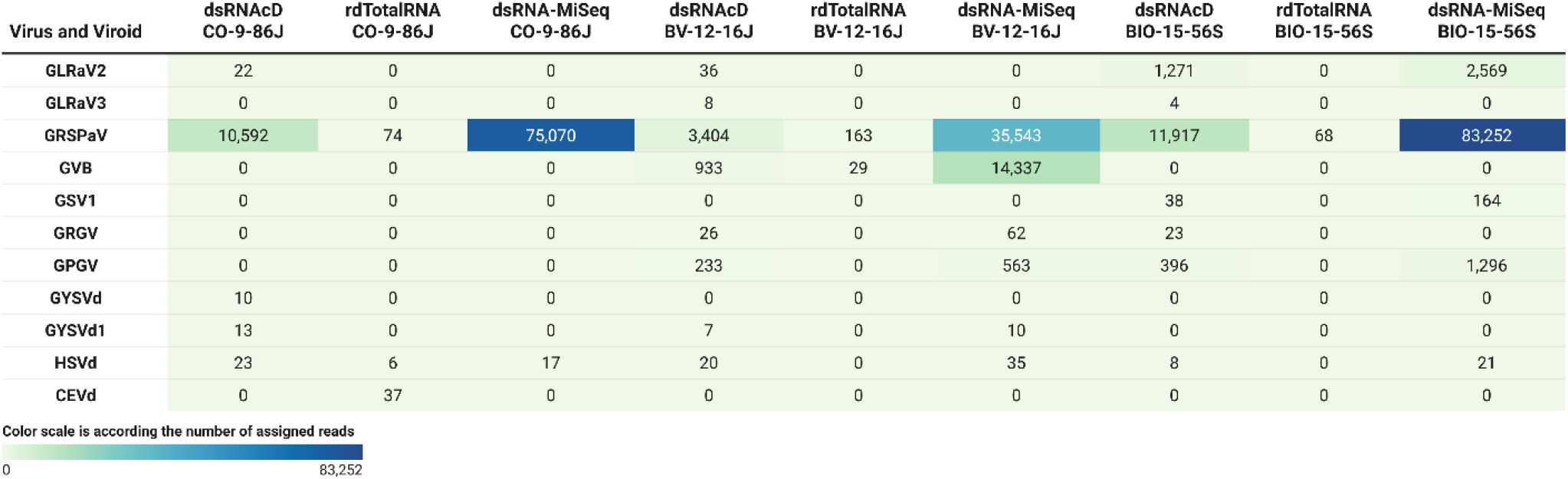
The number of assigned reads associated with grapevine viruses and viroids in different datasets. **GLRaV2**: grapevine leafroll-associated virus 2; **GLRaV3**: grapevine leafroll-associated virus 3; **GRSPaV**: grapevine rupestris stem pitting-associated virus; **GVB**: grapevine virus B; **GSV1**: grapevine Syrah virus 1; **GRGV**: grapevine red globe virus; **GPGV**: grapevine pinot gris virus; **GYSVd**: grapevine yellow speckle viroid; **GYSVd1**: grapevine yellow speckle viroid 1; **HSVd**: hop stunt viroid; **CEVd**: citrus exocortis viroid. (The heat map table was generated in https://www.datawrapper.de)

Cent&Rec analysis workflow was also used to compare viroid detection using dsRNA and total RNA. One unexpected viroid species, *Citrus exocortis viroid* (CEVd), was detected only in rdTotalRNA sequencing of CO-9-86J sample. However, the absence of CEVd in the dsRNAcD and dsRNA-MiSeq sequencing results raised a suspicion that these reads assigned to CEVd in the rdTotalRNA dataset might be reads from the host or other microorganisms. Accordingly, CEVd assigned reads were first extracted from the rdTotalRNA dataset of the CO-9-86J sample. Then blast alignment against ViroidDB and NCBI nt databases showed all CEVd-assigned reads were related to grapevines (*Vitis* spp.) (Supplementary file S5). According to the subject ID, the sequence of CEVd (FJ751964.1) in ViroidDB that was assigned to our sequences was retrieved from the NCBI website. Based on a blast against the NCBI nt database, the sequence of CEVd in ViroidDB showed 99.46% similarity to the sequences of grapevines (*Vitis* spp.), which indicated that the sequence was mistakenly annotated as CEVd in the ViroidDB database. Since the number of plant-originated reads in dsRNAcD datasets were lower than rdTotalRNA datasets, dsRNAs sequencing were more reliable than rdTotalRNA sequencing for viroid detection, even with the use of a viral-like entity dedicated and specialized database.

### Data analysis strategy 2: DIAMOND and MEGAN (DIA&MEG)

In addition to Cent&Rec, for the detection of viruses, we also used DIA&MEG workflow. Using DIAMOND, sequenced reads were aligned against the protein database (NCBI-nr), and then MEGAN processed the alignments to bin them based on taxonomic and functional information. Because the viroid genomes does not encode proteins, this strategy can only be used to detect viruses. The viruses detected in the dsRNAcD and dsRNA-MiSeq datasets through Cent&Rec were also detected by DIA&MEG (Figure. 5). Besides the results presented in figure 5, we also analyzed20 different datasets of dsRNA samples that were sequenced by dsRNAcD and dsRNA-MiSeq to verify the accuracy of the Cent&Rec workflow compared with the DIA&MEG workflow (Supplementary Folder 2).

**Figure 5.**
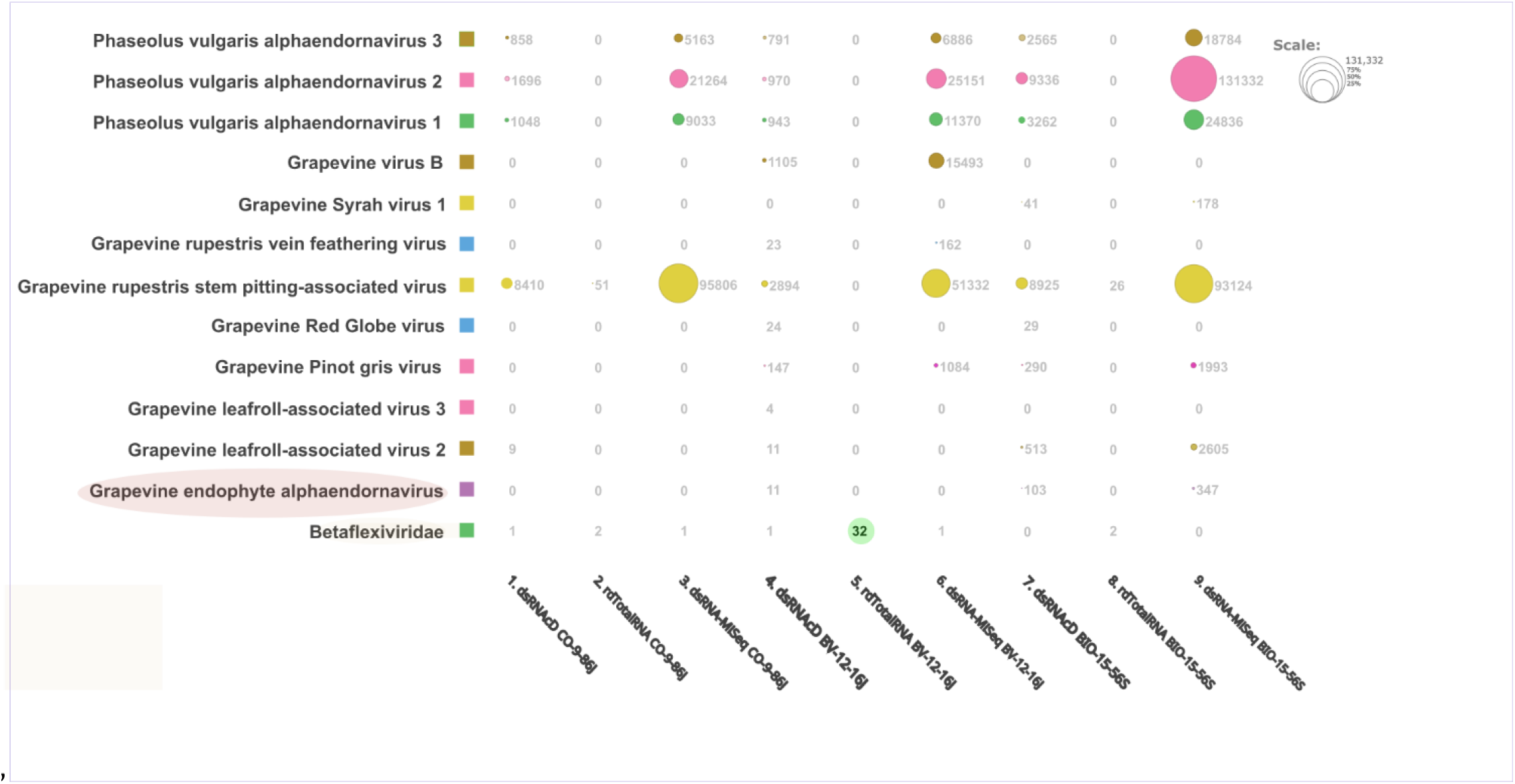
Comparative visualization of DIAMOND results by MEGAN Software. Multiple datasets (1. Nanopore dsRNAcD sequencing, 2. Nanopore rdTotalRNA sequencing, and 3. dsRNA-MiSeq sequencing) were uploaded to MEGAN and the relative abundance of reads in each sample was extracted.

The results from the DIA&MEG workflow were generally similar to those from Cent&Rec workflow except for GRSPaV, GVB and GEEV detection. GRSPaV was detected only in two of the three rdTotalRNA datasets when using DIA&MEG with default parameters, whereas it was detected in all three rdTotalRNA datasets when using Cent&Rec workflow (Figure 5). In the rdTotalRNA dataset of the BV-12-16J sample, GVBv and GRSPaV (from the Betaflexiviridae family) were detected by Cent&Rec, but not by DIA&MEG. However, 32 reads from the rdTotalRNA dataset of BV-12-16J sample were taxonomically assigned at the family level to Betaflexiviridae (Figure 6A). This result is explained by default threshold options in MEGAN software. Consequently, when two threshold options, “Min Support Percent” and “Min Support,” were turned off (=0), the taxonomic assignment of these 32 reads was done at the species level and these reads were assigned to GRSPaV and GVB (Figure 6B). Also, there were very few viral reads (34 reads) in the rdTotalRNA dataset of the BV-12-16J sample, even after removing host reads. The low abundance of viral reads in this dataset is a risk for accurate taxonomy classification. In contrast, both viruses, GRSPaV and GVB were binned taxonomically in dsRNAcD and dsRNA-MiSeq datasets correctly, even when using default threshold options in MEGAN software.

**Figure 6.**
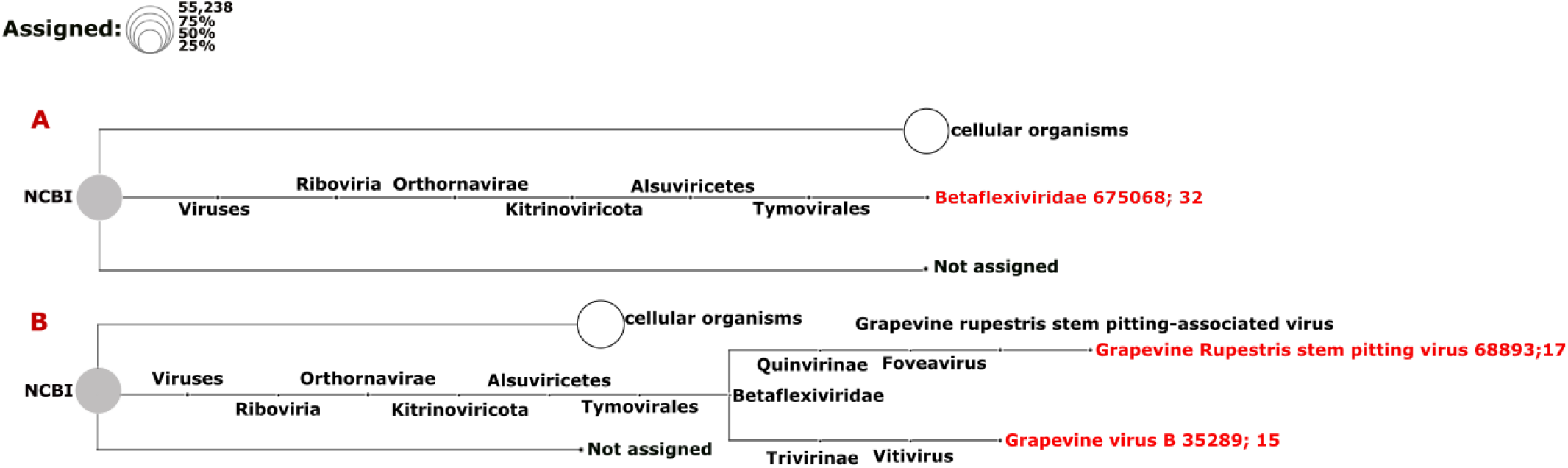
Assigning of viral reads to taxa using different thresholds in the rdTotalRNA dataset of the BV-12-16J sample. A. Taxonomic binning was done with default parameters of meganization option of MEGAN software. B. Two thresholds, “Min Support Percent” and “Min Support,” were turned off (=0) when meganization were done. There are two numbers following the virus name: a NCBI taxonomy ID followed by a number that indicates the number of assigned reads

A difference between dsRNA and total RNA datasets was also observed in the detection of GEEV. GEEV reads were not detected in the rdTotalRNA dataset for BIO-15-56S sample, but they were found in the dsRNAcD and dsRNA-MiSeq datasets (Figure 5). In addition, GEEV wasn’t detected in BIO-15-56S sample using Cent&Rec workflow, suggesting that the Cent&Rec results should be taken with caution. Since GEEV was detected only using the DIA&MEG workflow, we performed blast annotation against NCBI nt and nr databases to ensure GEEV assigned reads were not false positives. The blast results confirmed the presence of GEEV in the BIO-15-56S sample, which were in agreement with the results obtained by DIA&MEG (Supplementary file S6).

## Discussion

The aim of this study was to develop and evaluate two nanopore sequencing strategies for detecting viruses and viroids in mixed-infected grapevine samples: direct-RNA sequencing and direct-cDNA sequencing. We tried to answer two main questions relative to grapevine virus and viroids diagnostics. First, what is the best starting material for grapevine virus and viroid detection through single molecule nanopore sequencing, total RNA or dsRNA? Second, how cost-effective can nanopore-based diagnostic tool for grapevine virus and viroid detection be in comparison with MiSeq sequencing as a standard? In our study not only because viral reads were short (mean average was approximately 800bp), but also the number of viral reads was insufficient for de novo assembly. Therefore, two different taxonomical classification workflows based on protein and nucleotide homology, Cent&Rec and DIA&MEG, were evaluated for the detection of grapevine viruses and viroids.

Until now nanopore dsRNA sequencing has been used to identify or characterize viruses in single virus infections (45, 46). In fact, when it comes to identify and characterize the virome using nanopore sequencing technology, total RNA extraction is the common method used in the published literature. Several limitations have slowed the progress of using dsRNA for viral diagnostics, including labor-intensive dsRNA extraction methods and the notion that dsRNA may not be produced by some viruses. Our results showed preparation of a dsRNAcD sequecing library took more time than the preparation of a rdTotalRNA sequencing library. However, more samples were sequenced simultaneously on one flow cell (23 samples) through dsRNAcD sequencing, and the sequencing price per sample was three times lower (Can$103) than rdTotalRNA sequencing (Can$350). As part of the library preparation protocol for rdTotalRNA sequencing, two additional enzymatic steps, 1. rRNA-depletion and 2. polyadenylation, must also be used to remove unwanted rRNAs from samples and to capture non-poly A tailed viruses, which increased the sequencing price per sample. In contrast, by using dsRNA as the starting material and random primers for cDNA synthesis, these enzymatic steps were not needed, and the amount of unwanted sequences in dsRNAcD datasets was much lower than rdTotalRNA datasets. It remained to be determined how efficient these sequencing protocols are in terms of viral read yield, virus identification accuracy, and simplicity of bioinformatics data analysis.

The viral read portions in dsRNA datasets were significantly higher than in rdTotalRNAs datasets. For example, in the dsRNAcD dataset of CO-9-86J, 89% of assigned reads were related to viruses, while in the rdTotalRNA dataset only 6% of assigned reads were related to viruses. As well as failing to detect all viruses present in our samples, rdTotalRNA sequencing presented three other issues during data analysis. First, plant-derived reads were incorrectly assigned to Citrus exocortis viroid (CEVd) in the CO-9-86J sample. This issue was also reported in a previous study, where a plant sequence was misannotated as CEVd (72). In contrast, we did not observe this misannotation in our dsRNAcD and dsRNA-MiSeq datasets of the same sample. Because dsRNA datasets contain a smaller proportion of plant-derived sequences, dsRNA is a more reliable starting material for detecting viruses and viroids through taxonomical classification, thus decreasing the likelihood of false positive results.

Second, there was a problem related to taxonomic classification of viruses in the rdTotalRNA dataset of BV-12-16J. Since the abundance of GRSPaV and GVB reads in rdTotalRNA dataset was low, taxonomic classification at the species level weren’t possible when using default setting of MEGAN. Taxonomy classification of long reads, specifically viral reads, can be affected by a number of factors including viral load and mutation rate. Indeed, species with low abundance can be discarded according to threshold options in the taxonomy classifier software. In order to apply thresholds effectively and avoid discarding low abundance species, it is important to take into consideration the type of study, read depth, and classifier software used (73). Furthermore, most taxonomic classifiers are designed to classify bacteria or microorganisms rather than viral reads. The high mutation rate of viruses should be considered, and consequently the software’s mismatch threshold may need to be adjusted (74). In contrast, both dsRNAcD and dsRNA-MiSeq yielded enough viral read counts for accurate detection of both GRSPaV and GVB when using the default MEGAN thresholds.

Third, the detection of GEEV was also affected by the type of starting materials used for sequencing and the bioinformatics workflow used. The dsRNAcD and dsRNA-MiSeq datasets of a sample contained viral reads related to GEEV, while the rdTotalRNA dataset of the same sample did not. In addition, Cent&Rec workflow wasn’t able to detect GEEV in all datasets of some samples. In dsRNA datasets, although the proportion viral reads assigned reads was high, there was a substantial amount of unassigned reads as well. While reverse transcription artifacts could explain the high number of unassigned reads (75) in dsRNA sequencing datasets, some of these unassigned reads may have been related to novel virus strains or species. We conducted further curation by BLASTn and BLASTx in order to confirm that the detected GEEV was not a false positive. The results indicated that the sequence similarity between detected GEEV and the NCBI record (YP_007003829.1) was at the protein level, not at the nucleotide level, suggesting that different strains or haplotypes of GEEV may exist in that grapevine sample. However, further studies will be needed to confirm the genetic diversity within GEEV species which this was beyond the scope of this study. According to Al Rwahnih et al., dsRNA sequencing produced a grater number of unique cDNA sequences than total RNA sequencing, thus offering a greater possibility of discovering new viruses. Indeed, several numbers of novel grapevine viruses including grapevine Syrah virus-1, grapevine red blotch virus etc., were discovered through dsRNA sequencing (10, 76). Therefore, by using dsRNA, not only known viruses and viroids can be detected, but the potential for discovering new virus strains or species is increased.

However, the remaining question is how cost-effective nanopore dsRNA sequencing would be in comparison to Illumina MiSeq sequencing? To determine which sequencing technology would be most appropriate for diagnostic laboratories and disease management projects, we compared cost-effectiveness of both MiSeq and nanopore sequencing. We compared them in terms of library preparation time, ease of use, sequencing price, and data analysis requirements. In general, the cost-effectiveness of using Illumina sequencing in diagnostic lab discourage the use of this technology for day-to-day diagnostic activities. Indeed, the cost of the sequencer and the minimum number of samples required (50 to 60 samples) to be cost-effective make Illumina sequencing less suitable for small diagnostic labs. Nanopore sequencing is an alternative to MiSeq sequencing in this situation. In our study, the estimated cost for nanopore dsRNA sequencing was Can$103 per sample whereas Gaafar and Ziebell, 2020 (49) estimated the cost for virus detection by MiSeq sequencing to Can$412 per sample, which indicates nanopore dsRNA sequencing is four times cheaper than dsRNA-MiSeq sequencing. In addition, a nanopore dsRNAcD sequencing library can be prepared and sequenced in 37.58 h, whereas a dsRNA-MiSeq library takes nearly 88.75 h to prepare and sequence (Table 2). Our analysis showed that the detected virus list of nanopore dsRNAcD sequencing using both proposed data analysis workflows, DIA&MEG and Cent&Rec was similar to the dsRNA-MiSeq results in all samples. While there wasn’t any difficulty choosing a pipeline for analyzing MiSeq data, there were some challenges in analyzing nanopore sequencing data. Similar to the situation described by Amoia et al. 2022 (45), our dsRNAcD libraries provided relatively low quality, and low quantities of viral reads to meet the requirements of long read based de novo assembler software. Thus, these software was not able to assemble long and numerous viral contigs. Most de novo assemblers are designed for reconstructing complete microbial and near-complete eukaryotic genomes. Therefore, we had to analyze our raw data by taxonomy classifier tools. Also, because nanopore sequencing raw data trimming, filtering, and error correction take more time, virus detection was slower than MiSeq data analysis. A further step in data pre-processing that we performed was the subtraction of host-originating reads from trimmed and error corrected datasets with the goal of increasing viral read counts and improving the taxonomic classification of viruses. Despite the fact that DIAMOND+MEGAN workflow takes a long time to run for the analysis of large datasets, such as environmental metagenomic datasets (66, 67), it took a short time (around two hours) to run on our datasets. This workflow allows the user a wide range of options, including displaying taxonomic tree of detected viruses, extracting functional information from assigned reads, compares different samples, and analyzes and compares short read and long read sequencing datasets. However, the Cent&Rec workflow was faster than DIAMOND+MEGAN in terms of running time (around 20 to 30 minutes), and this workflow was able to detect viroids contrary to DIAMOND+MEGAN workflow. Our study demonstrated that dsRNAcD sequencing can effectively compete with MiSeq sequencing for the detection of viruses and viroids.

## Conclusion

The purpose of this study was to examine the ability of nanopore direct cDNA and direct RNA sequencing technologies to detect simultaneously grapevine viruses compare to Illumina MiSeq sequencing. According to our results, dsRNA as the starting material for library preparation is more reliable than total RNA in terms of identifying grapevine viruses and viroids. For all samples, the dsRNA sequencing results were similar to those from dsRNA-MiSeq sequencing. In addition, the rRNA depletion step did not improve grapevine viruses detection from total RNA libraries despite the increase in the cost per sample. In contrast, when dsRNA was sequenced using direct cDNA sequencing kit, not only more samples (23 samples) were multiplexed and sequenced simultaneously on one flow cell, but also the total cost of sequencing has fallen significantly (Can $103/sample). However, there is a need to improve and optimize the current dsRNA purification protocol to gain efficiency in time and effort. In conclusion, the study demonstrated that the dsRNAcD sequencing can be a routine and affordable diagnostic tool for detecting plant viruses.

## Acknowledgment

We are grateful to Dong Xu from the Saint-Jean-sur-Richelieu Research and Development Center (CRDH) of Agriculture and Agri-Food Canada and Sylvain Lerat from Département de Biologie, Centre SÈVE, Université de Sherbrooke for their support and assistance with plant sampling and processing. Additionally, We would also like to thank Joël Lafond-Lapalme and Pierre-Yves Véronneau for technical and bioinformatic assistance, as well as Sarah Drury for reviewing the paper.

